# Structural investigation of ACE2 dependent disassembly of the trimeric SARS-CoV-2 Spike glycoprotein

**DOI:** 10.1101/2020.10.12.336016

**Authors:** Dongchun Ni, Kelvin Lau, Frank Lehmann, Andri Fränkl, David Hacker, Florence Pojer, Henning Stahlberg

**Author notes:** **Corresponding author**: Henning Stahlberg, Center for Cellular Imaging and NanoAnalytics (C-CINA) Biozentrum, University of Basel, WRO-1058, Mattenstrasse 26 CH-4058 Basel, Switzerland.

## Abstract

The human membrane protein Angiotensin-converting enzyme 2 (hACE2) acts as the main receptor for host cells invasion of the new coronavirus SARS-CoV-2. The viral surface glycoprotein Spike binds to hACE2, which triggers virus entry into cells. As of today, the role of hACE2 for virus fusion is not well understood. Blocking the transition of Spike from its prefusion to post-fusion state might be a strategy to prevent or treat COVID-19. Here we report a single particle cryo-electron microscopy analysis of SARS-CoV-2 trimeric Spike in presence of the human ACE2 ectodomain. The binding of purified hACE2 ectodomain to Spike induces the disassembly of the trimeric form of Spike and a structural rearrangement of its S1 domain to form a stable, monomeric complex with hACE2. This observed hACE2 dependent dissociation of the Spike trimer suggests a mechanism for the therapeutic role of recombinant soluble hACE2 for treatment of COVID-19.

## Introduction

Coronavirus is a family of single-stranded RNA viruses, many of which can infect animals and humans (MacLachlan and Dubovi, 2017; Monto, 1984). The symptoms of coronavirus-related diseases can be mild and mainly occur in respiratory tract. For example, roughly 15%-30% cases of the common cold are caused by human coronaviruses (Mesel-Lemoine et al., 2012). A coronavirus infection sometimes can develop serious illnesses, such as SARS (severe acute respiratory syndrome), MERS (Middle East respiratory syndrome) and also the current pandemic COVID-19 (coronavirus disease 2019) (Tang et al., 2020). SARS-CoV is a beta-coronavirus that caused a pandemic in 2002. SARS-CoV-2 is a novel coronavirus that is genetically similar to the previous SARS-CoV. SARS-CoV-2 causes the ongoing pandemic COVID-19 and has been spreading globally since the first quarter of this year (Ciotti et al., 2020). The symptoms of COVID-19 vary from person to person. In some cases, the illness is very serious, in particular for the elderly (Pascarella et al., 2020). As of today, no specific antiviral drugs were approved for use against COVID-19 and vaccine development is still at the phase of clinical testing.

Cryogenic electron microscopy (Cryo-EM) is a technique for structure determination of biomacromolecules, which has been particularly successful for studying high molecular-weight proteins. Cryo-EM does not require crystallization of the target protein. COVID-19 related protein structures have been widely investigated since the spread of SARS-CoV-2 in this year, using cryo-EM single particle analysis (SPA). The protein nicknamed Spike is with its 180kDa monomeric molecular weight the largest viral surface protein of SARS-CoV-2. It consists of two domains S1 and S2 that are connected by a short linker. Spike forms stable trimers on the virus surface that are attached to the virus membrane. This Spike trimer is the key molecule for host cells receptor binding and invasion of the host cells. The cryo-EM structure of the entire Spike homotrimer was determined recently, showing a mushroom sharped overall architecture (Walls et al., 2020; Wrapp et al., 2020a). As also for the SARS-CoV, the viral fusion bridge from SARS-CoV-2 to the host cell is formed by Spike and the ectodomain of the human Angiotensin-converting enzyme 2 (hACE2), which is the virus receptor on the host cell that triggers virus entry. *In vitro* studies have shown that the Spike receptor binding domains (RBDs) from SARS-CoV as well as SARS-CoV-2 can both bind to the ectodomain of hACE2 with comparable binding affinities in low nanomolar levels (Lan et al., 2020a). However, the new SARS-CoV-2 exhibits a more potent capacity of host cells adhesion, as well as a larger virus-entry efficiency than other beta-coronaviruses (Shang et al., 2020).

The membrane-attached hACE2 is known to be the key molecule for the infection by several viruses, including SARS-CoV, Human coronavirus NL63 (HCoV-NL63) and SARS-CoV-2. The infection process primarily involves virus adhesion and fusion (Bao et al., 2020; Fehr and Perlman, 2015; Kuba et al., 2005; Sia et al., 2020). Interestingly, hACE2 may not only serve as a drug target to prevent SARS-Co-2 infection, but hACE2 itself may also be considered as a potential therapeutic drug candidate for the usage against COVID-19 or other beta-coronavirus related diseases. The clinical-grade soluble form of hACE2 has been reported to be a potential novel therapeutic approach for reducing the infection of SARS-CoV-2 (Monteil et al., 2020) by preventing the viral Spike from interacting with other hACE2 present on human cells. Recently, researchers have also characterized the entire architecture of the inactivated authentic virions from SARS-CoV-2 using cryo-electron tomography, observing that post-fusion S2 trimers are distributed on the surface of SARS-CoV-2 virions (Ke et al., 2020; Turonova et al., 2017). The exact role of hACE2 so far is not yet fully understood in terms of its interaction with full-length Spike protein.

In this study, we present a cryo-electron microscopy (cryo-EM) study of the SARS-CoV-2 Spike protein in complex with hACE2. Our analysis reveals a monomeric complex of Spike S1 domain with hACE2, requiring a large structural rearrangement in S1 compared to its isolated structure. Our data show that hACE2 binding induces a conformational change in Spike, leading to Spike trimer dissociation.

## Results

### Spike and hACE2 production and its complex assembly

The prefusion Spike 2P ectodomain was expressed in ExpiCHO cells and affinity purified via its twin Strep-tag. SDS-PAGE analysis showed the presence of pure full-length Spike protein, consisting of both, the S1 and S2 domains at the expected molecular weight of 180 kDa for the Spike monomer. (**Suppl. Fig. S1a**). The purified Spike sample in PBS buffer was imaged as negatively stained preparations by transmission electron microscopy (TEM). This revealed the expected trimeric shape, and 2D class averages of selected particles in negative stain TEM images showed the typical, mushroom-shaped particles (**Suppl. Fig. S2a.c**), in accordance with the expected structure of the SARS-CoV-2 Spike in the pre-fusion state.

Human ACE2 ectodomain was expressed in HEK293 cells and purified via a poly-histidine immobilized metal affinity chromatography (IMAC) with a Fastback Ni2+ column, followed by another anion exchange column (**Suppl. Fig. S1b**). For details, see Methods.

Purified Spike protein was mixed with excess hACE2 (molar ratio Spike:hACE2 of 1:5) and incubated for 12 hours at 4° C. Samples were prepared by negative staining and imaged by TEM. Unexpectedly, the observed particle features were largely different from the typical Spike trimers in shape. A 2D analysis of 5’854 picked negatively stained particles revealed in class averages that the majority of the particles were smaller in size and asymmetrical, compared to the non-incubated Spike trimeric samples (**Suppl. Fig. S2b,d**). This suggests that the prolonged incubation with hACE2 led to Spike trimer dissociation. A similar observation for SARS-CoV-ACE2 complex was recently reported by Song et al (Song et al., 2018). Due to particle heterogeneity, we decided to further purify the complex by size exclusion chromatography (SEC) and indeed the SEC profile showed three distinct peaks, called Peak1, Peak2 and Peak3 (**Suppl. Fig. S1c,d**).

By analyzing the 3 peaks by SDS gel and negative stain EM, we could clearly differentiate non-structured aggregates of full-length Spike and hACE2 in Peak1, that did not allow further structural analysis (data not shown), to the excess of unbound hACE2 in Peak3 (**Suppl. Fig. S1c,d**). The homogenous Peak2 that contained full-length Spike in complex with hACE2, was further analyzed by cryo-EM.

### hACE2 binding can induce disassembly of Spike homotrimer

Peak2 (Spike:hACE2 at molar ratio 1:5 after overnight incubation at 4°C) was vitrified and frozen grids were loaded into a Thermo Fisher Scientific (TFS) Titan Krios cryo-EM instrument, operated at 300kV acceleration voltage, and equipped with a Gatan Quantum-LS energy filter equipped with K2 direct electron detector **(Suppl. Fig. S3)**. 8’927 dose-fractionated images (i.e., movies) were recorded **(Suppl. Fig. S4**), from which ~1.7 million particles were extracted and subjected to image processing and 3D reconstruction. The final 3D reconstruction from 72’446 particles at 5.1Å overall resolution showed a density map corresponding to a single, monomeric Spike protein in complex with hACE2 (**Fig. 1a**). The map allowed docking with available structures for S1 and hACE2 taken from the previously reported structures (Spike PDB ID 6VYB and Spike RBD-ACE2 6M0J), revealing a structural rearrangement of the C-terminal domain (CTD) and N-terminal domain (NTD) of S1 compared to a monomer from that Spike structure in the RBD^up^ conformation. The interaction between the S1 RBD and hACE2 is in agreement with several other reported structures of the RBD-ACE2 complex (PDB ID 6M0J, 6VW1 or 6LZG). No additional density for S2 or a fragment of S2 was detected in the reconstruction.

**Figure 1.**
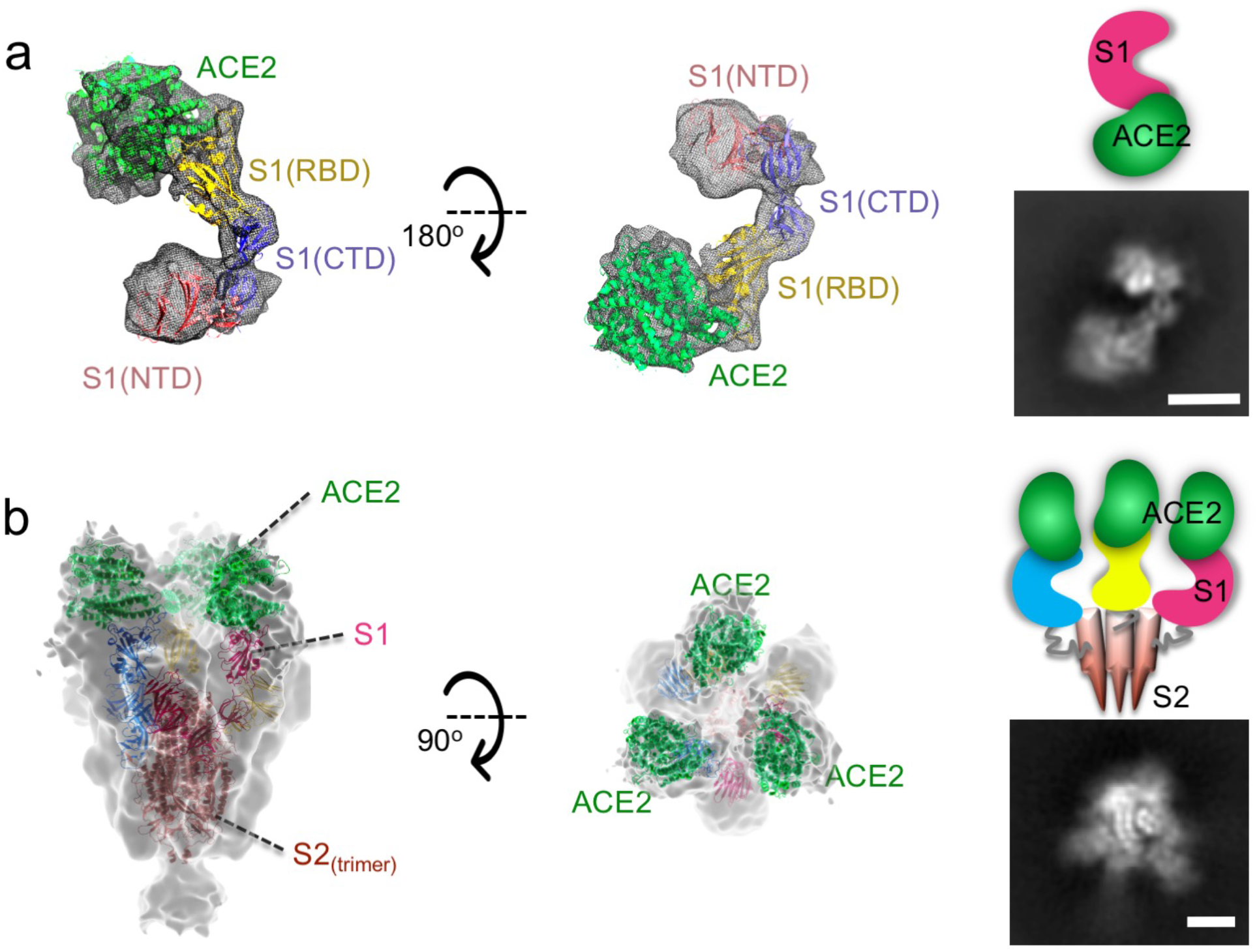
Cryo-EM maps of SARS-CoV-2 Spike-hACE2 complexes and fitted models. **a**. The 3D reconstruction of SARS-CoV-2 Spike and human ACE2 (mixed at a molar ratio 1:5) incubated for 12hrs and further purified by SEC shows a structure corresponding to monomeric Spike S1 protein in complex with hACE2. No density for S2 is observed. The N- and C-terminal domains of S1 had to be re-arranged to fit into the map. Left: Side-view, center: 90° rotated view. The structure is colored as follows: hACE2 ectodomain green, Spike S1-RBD yellow, Spike S1-CTD slate blue, Spike S1-NTD salmon. Right: The bottom image shows a representative 2D class average of the S1-hACE2 complex. The upper cartoon is its interpretation. **b**. 3D reconstruction of Spike and hACE2 (ratio 1:3) incubated for 3 hours shows a trimeric map allowing docking of three hACE2 molecules and three S1 and three S2 molecules, all forming a trimeric complex. The map is low-pass filtered to 9Å resolution for better interpretation. The complex is arranged in a molar ratio of 3:3 (SpikeS1-S2:hACE2). The image on the left indicates the host cell membrane as a cartoon. Center: 90° rotated view. Right: The bottom image shows a representative 2D class average of Spike-hACE2 (3:3) complex, here shown from the top as in the central panel. The upper cartoon is its interpretation (here shown from the side). Scale bar in class averages: 2 nm.

We tested a shorter incubation time by mixing Spike:hACE2 (molar ratio of 1:3) and let it incubate for 3 hours at 4°C, instead as overnight. No further SEC purification was performed. Subsequently, cryo-EM grids of this sample were prepared and subjected to cryo-EM analysis (**Suppl. Fig. S5)**. From 7’045 recorded movies, 615’348 particles were extracted and subjected to classification and 3D analysis. This revealed a small sub-set of 47’901 particles corresponding to the prefusion Spike trimer, which allowed a 3D reconstruction at 4.2Å overall resolution (no symmetry was applied), while some regions of the 3D map showed lower resolution, presumably due to increased flexibility of these areas (**Suppl. Fig. S6**). A resolution-limited map at 9Å resolution (**Fig. 1b**) allowed clear docking the models of Spike S1 and S2 and hACE2, which showed that the complex is composed of Spike and hACE2 in a molar ratio of 3:3 (Spike:hACE2). Three hACE2 molecules were observed to attach to the RBDs of Spike. All three RBDs were in the RBD^up^ conformation and slightly shifted away from the central trimer axis (**Fig. 1b**). A similar arrangement was also recently observed by Kirchdoerfer et al. (Kirchdoerfer et al., 2018), see also (Zhou et al., 2020).

### Structural comparison of Spike-hACE2 complexes

A detailed analysis of both obtained structures shows significant structural rearrangements in different forms of Spike-hACE2 complex. The resolution of our reconstructed EM maps was not sufficient for building atomic models, but allowed clear placement of available atomic structures for Spike (PDB ID: 6VYB) and ACE2-RBD (PDB ID: 6M0J) (Lan et al., 2020b; Walls et al., 2020). Comparison of the docked and fitted monomeric S1-hACE2 model to that of the trimeric Spike-hACE2 (3:3) showed that a ~30° rotation of the C-terminal and N-terminal sub-domains of Spike S1 were required to bring the Spike S1 protein into the monomeric arrangement with hACE2. After such re-arrangement, the domains of hACE2 and the RBDs of the S1 protein are in good agreement with a reported crystal structure (REF Lan et al.) (**Fig. 2b)**.

**Figure 2.**
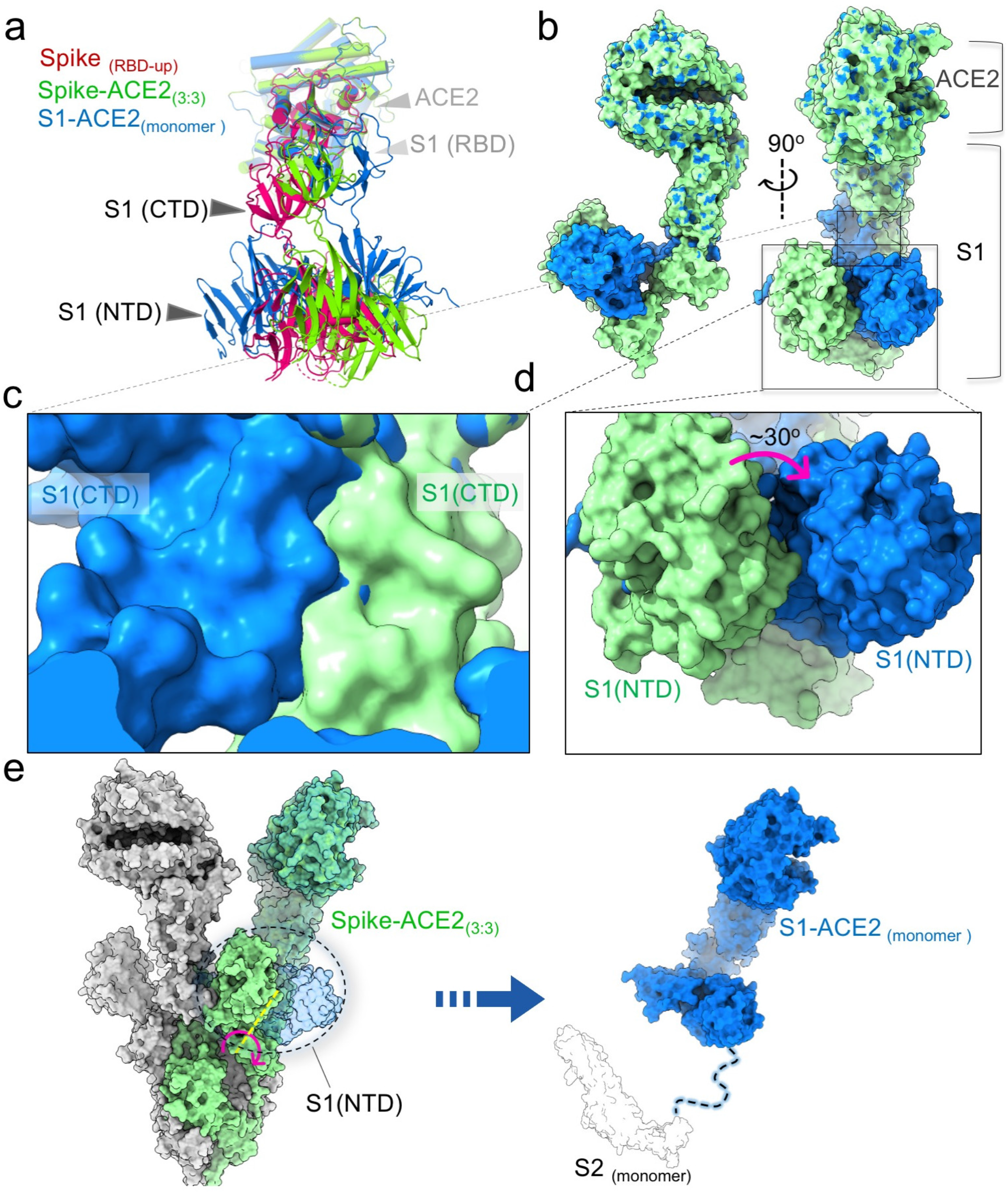
Conformational rearrangement of Spike-hACE2. **a**. Superposition of S1-hACE2 complex structures derived from different conformations. Structures are shown as a cartoon and colored as follows: monomeric Spike-hACE2 complex, blue; trimeric form of Spike-hACE2 complex, green; Spike one RBD-up (PDB ID: 6VYB), red. The RBDs have been superimposed. **b**. Structural comparison of S1-hACE2 regions. The movement is presented as two colors: SARS-CoV2 trimeric Spike-hACE2 complex (green) and S1-hACE2 (blue) **c,d**. The close-up views show the proposed local structural rearrangements. **e**. Superposition of S1-hACE2(1:1) with the trimeric form of Spike-hACE2 complex. The RBDs have been superimposed. The rearranged structure of Spike-hACE2 is no longer compatible with formation of trimers so that it dissociates. The right panel is a predicted structure of the dissociated Spike-hACE2 monomer.

When comparing the docked model of the monomeric S1-hACE2 complex with that of the trimeric Spike-hACE2 complex (**Fig. 2e**), a considerable number of stearic clashes at the interface between Spike S1 (CTD) and its neighboring region from the S2 polypeptide chain was obvious. The docked monomeric S1-hACE2 complex is further structurally incompatible with the observed trimeric arrangement.

## Discussion

The stoichiometric ratio of the complex of Spike:hACE2 on the host cell upon virus entry is not well established. Nevertheless, one hACE2 molecule per Spike trimer is likely sufficient for binding and initializing virus fusion with the host cell (Song et al., 2018). Even though the structure of a post-fusion S2 trimer has recently been determined (Cai et al., 2020), it is not clear how membrane fusion during virus entry is coordinated upon release of the S1-hACE2 caps (**Fig. 3**). We here report the cryo-EM structure of a stable monomeric S1 Spike-hACE2 (1:1) complex. Even though size-wise it would have been detectable, our particle classification did not reveal any particle class corresponding to an isolated S1 fragment in addition to the observed S1 Spike-hACE2 (1:1) particles (**Suppl. Fig. S3c, S4)**. Knowledge of the mechanism how the S1 fragment might be detached from the hACE2 receptors after virus entry would be relevant for understanding its mechanism of infection and pathogenicity. The fact that we did not observe any free S1 fragments suggests that the S1-hACE2 complex is rather stable, at least under our *in vitro* conditions.

**Figure 3.**
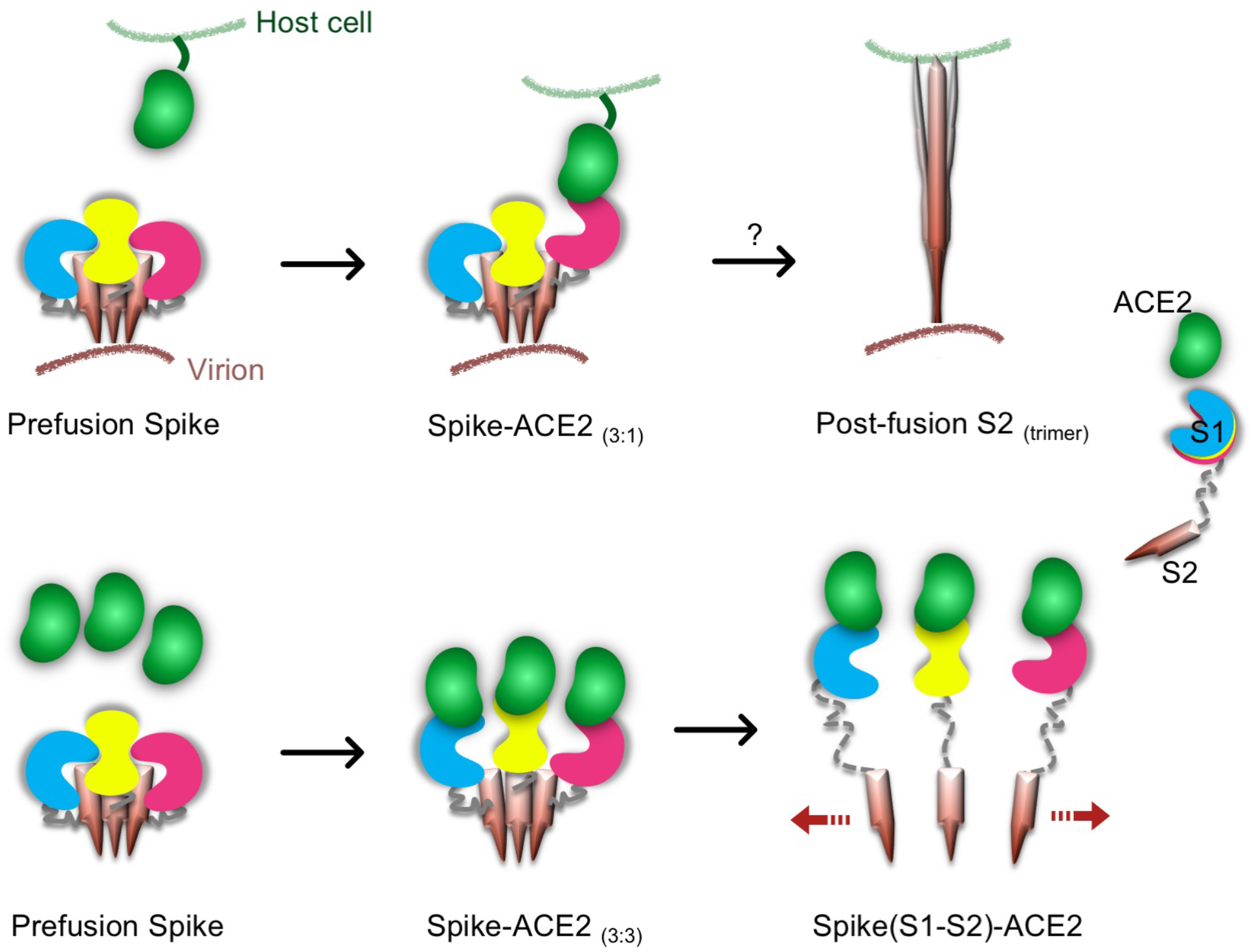
Proposed models for Spike-hACE2 complex formation and structural rearrangement. The upper row shows a possible pathway leading to a conformational change of the trimeric SARS-CoV2 Spike. In this model, one hACE2 molecule binds to one Spike S1 monomer and induces the conformational changes in the trimeric Spike. Subsequently, a post-fusion S2 trimer is formed. The lower row shows a novel proposed pathway leading to Spike trimer disassembly by hACE2. In presence of a high concentration of hACE2 molecules, a Spike-hACE2 (3:3) complex is formed. Structural clashes between the three Spike-hACE2 elements lead to their dissociation. This induces the formation of monomeric Spike-hACE2 complexes.

Secondly, the S2 domain was not detected in the obtained structure of the Spike-hACE2 monomeric complex (1:1), even though the SDS-PAGE analysis showed that S2 was present as full length in the sample (**Suppl. Fig. S1d**). The S2 domain is expected to be connected to the S1 domain via a short loop between S1 and S2, where a Furin protease cleavage site is expected (Belouzard et al., 2009; Haan et al., 2004; Hoffmann et al., 2020). However, in the absence of stable trimers, the loop between S1 and S2 is likely very flexible, possibly making the S2 domain undetectable by cryo-EM maps. Our cryo-EM analysis that didn’t show the S2 domain in the averaged 3D reconstruction therefore likely failed to align the S2 domains either due to their flexibility, or due to a denaturation of S2 during sample preparation.

An early study presented a potential dose-dependent inhibition of SARS-CoV-2 infection by a recombinant soluble form of hACE2 (Monteil et al., 2020). The mechanism, how the soluble forms of hACE2 would be able to neutralize the virus, is not known. One possible mechanism could be a direct competition between the soluble hACE2 and the host cell hACE2 receptor, so that Spike proteins saturated with soluble hACE2 domains render them unable to interact with host cell hACE2. Here, however, we report that the soluble forms of hACE2 induce the opening and disassembly of the trimeric Spike structure to create the stable Spike S1-hACE2 complex (**Fig. 3**). We propose a mechanism by which the formation of the Spike-hACE2 (3:3) complex induces a high structural flexibility in the Spike trimer, allowing a conformational re-arrangement of the S1 C- and N-terminal domains when interacting with hACE2. In consequence, the new S1-hACE2 complex is incompatible with a trimeric arrangement, causing the dissociation of the trimeric complex (**Fig. 3**).

This hypothesis is supported by the recent manuscript deposited in bioRxiv.org, which describes a similar effect triggered by engineered DARPin molecules (Walser et al., 2020). Therefore, we suppose that the soluble forms of hACE2 may not only block the infection and replication of SARS-CoV-2, but also destroy the trimeric Spike adaptors that are responsible for viral host membrane fusion. This mechanism suggests a novel therapeutic strategy for the treatment of COVID-19, by adding soluble hACE2 to dissociate the Spike trimer of approaching viruses.

## Methods

### Protein production and purification

#### Spike protein

The gene for the prefusion ectodomain of the SARS-CoV2 Spike 2-P protein (the prefusion stabilized construct (2P) that includes the putative furin cleavage site mutated; the plasmid was a generous gift from Prof. Jason McLellan, University of Texas, Austin (Wrapp et al., 2020b)) was transiently transfected into suspension-adapted ExpiCHO cells (Thermo Fisher) with PEI MAX (Polysciences) in ProCHO5 medium (Lonza). After 1 h, dimethyl sulfoxide (DMSO; AppliChem) was added to 2% (v/v). Incubation with agitation was performed at 31°C and 4.5% CO_2_ for 5 days. The transparent supernatant was passed over a Strep-Tactin column (IBA Lifesciences) and bound protein was eluted with PBS buffer + 2.5 mM desthiobiotin. The elute was dialyzed against PBS buffer. The average protein yield for Spike 2P was 15 mg/L culture.

#### hACE2 protein

The gene for the secreted expression of hACE2-Fc-His was a generous gift from Prof. Jason McLellan. The construct was transiently transfected into HEK293 cells (Thermo Fisher) with PEI MAX (Polysciences) in EX-CELL 293 serum-free medium (Sigma), supplemented with 3.75 mM valproic acid. Incubation with agitation was performed at 37°C and 4.5% CO_2_ for 8 days. The clear supernatant was purified as follows via MabSelect Protein A resin (Cytiva), cleavage of the Fc-His tag by overnight treatment with 3C protease, removal of the cleaved tag and 3C protease by NiNTA followed by an anion exchange column. The purified protein was finally dialyzed into PBS buffer. The average yield for cleaved hACE2 was 13 mg/L culture.

### Negative stain TEM imaging

For all protein samples, 3ul of sample solution was applied to glow-discharged carbon-coated Cu-grids and incubated for 1min. The free liquid was removed by blotting with filter paper and the grids were stained for 25s with 2% uranyl acetate solution. The negatively stained preparations were imaged with a Tecnai G2 Spirit TEM, operated at 120kV at a magnification of 135kx. Image were recorded with a Veleta CCD camera (EMSIS GmbH, Münster, Germany). The 2D analysis of the negative stain TEM images was performed with the program cisTEM1.0 (Grant et al., 2018).

### Cryo-EM imaging

Two different preparations of cryo-EM grids were performed. Purified Spike and hACE2 proteins were mixed at the molar ratio of 1:5 and incubated for 12 hours at 4°C. After that, the sample was subjected to size exclusion chromatography (SEC) with a Superose 6 increase (10/300) column, and the fractions from Peak2 were pooled and concentrated in 100 kDa centrifugal concentrators (Millipore).

Alternatively, purified Spike and hACE2 were mixed at the molar ratio of 1:3 (Spike:hACE2) an incubated for 3 hours at 4°c, without further purification via sec.

For both samples, the concentration was adjusted to 0.5mg/ml. Cryo-EM grids were prepared with a Vitrobot Mark IV (Thermo Fisher), using a temperature of 4°C and 100% humidity. 4 μL of sample was applied onto glow-discharged Quantifoil holey carbon grids (R2.0/2.0, 200 mesh, copper) and blotted for 2.0-3.0s. The grids flash-frozen in a liquid ethane, cooled by liquid nitrogen.

For both samples, dose-fractionated images (i.e., movies) were recorded with a Titan Krios (Thermo Fisher), operated at 300kV, and equipped with a Gatan Quantum-LS energy filter (20 eV zero-loss energy filtration) followed by a Gatan K2 Summit direct electron detector. The data collection statistics is presented in **Table 1**. Images were recorded in counting mode, at a magnification yielding a physical pixel size of 0.82Å at the sample level. Images were automatically recorded with the SerialEM program (Mastronarde, 2003) at a defocus range of −0.8 ~ −2.5um. All the movies were gain-normalized, aligned, dose weighted and averaged with the program of MotionCor2 (Zheng et al., 2017) within FOCUS (Biyani et al., 2017), which also was used to sort images and reject images of insufficient quality. The pre-processed micrographs were imported into cryoSPARC V2 (Punjani et al., 2017).

**Table 1:**
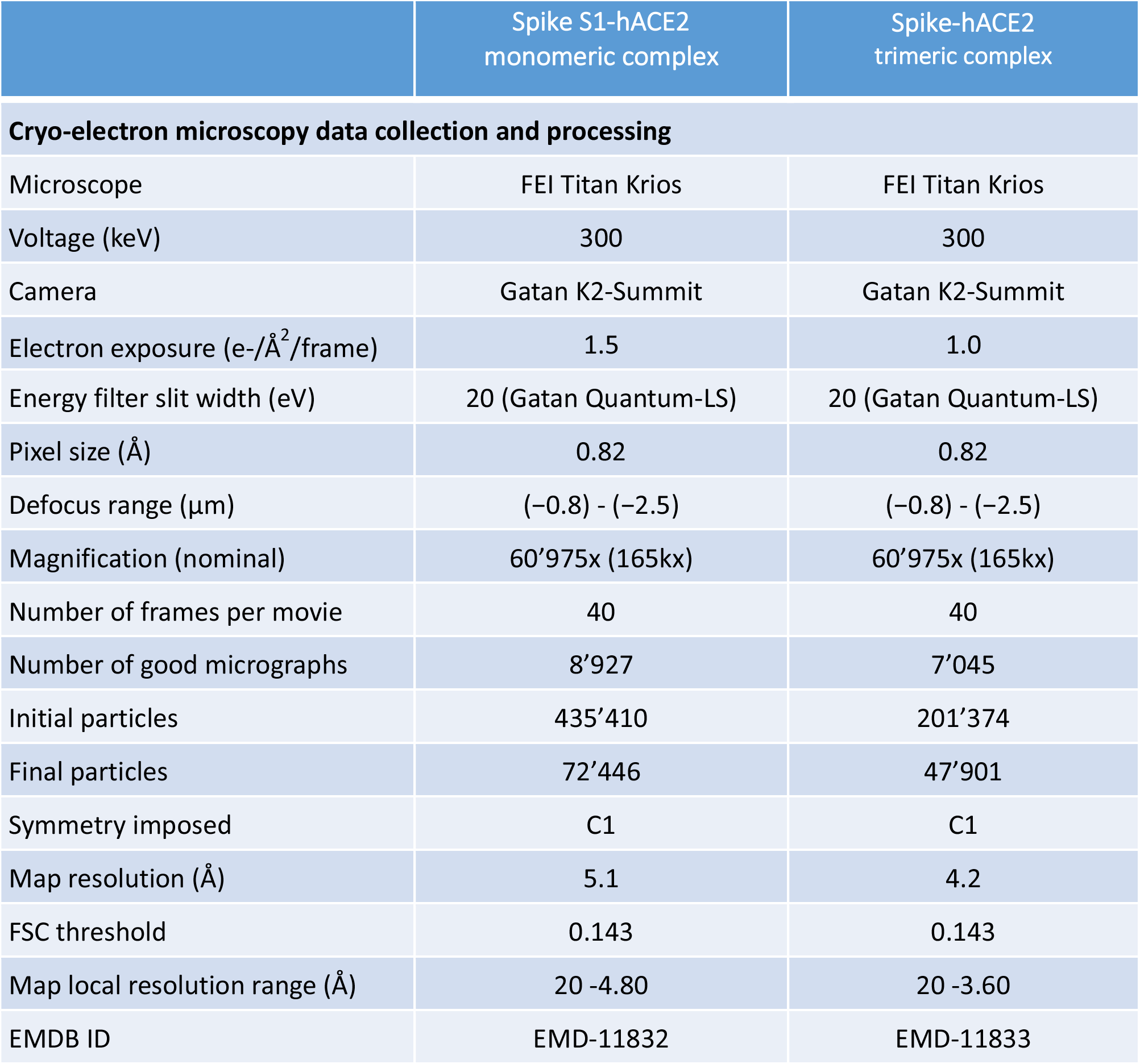
EM data collection statistics.

### Image processing

Data processing of the monomeric Spike-hACE2 complex was conducted in CryoSPARC V2 (Punjani et al., 2017). The defocus and contrast transfer function (CTF) values were estimated on 8’927 micrographs and 7’927 good images were selected. 1’685’202 particles were auto-picked by blob picking, followed later by another round of template picking. Particle clearing was performed by two rounds of 2D classifications, resulting in a particles stack of 435’410 particles. 3D references were generated using ab-initio reconstruction (CryoSPARC V2) and followed by two rounds of 3D hetero-refinements. The final particle set contained 72’446 particles, leading to a 3D map at 5.1Å overall resolution, as estimated by Fourier Shell Correlation (FSC) using the 0.143 cutoff criterion.

For the trimeric Spike-hACE2 complex, image processing was performed similarly. The final reconstruction produced a 3D map at 4.2Å overall resolution.

### Model interpretation

Protein models were generated from reported structures (Spike: PDB ID 6VYB; ACE2-RBD: PDB ID 6M0J) (Lan et al., 2020b; Walls et al., 2020). For the S1-hACE2 structure, the model was manually docked into the EM density with the program Chimera (Pettersen et al., 2004) and further refined using rigid-body fitting in COOT (Emsley et al., 2010). For the Spike-hACE2 trimer, the density corresponding to hACE2 was relatively weak, so that low-pass filtration to 9Å resolution was applied to the map before proceeding with the docking of hACE2 as described above.

## Figure preparation

Figures were created using the software PyMOL (PyMOL Molecular Graphics System, DeLano Scientific), Chimera and ChimeraX (Goddard et al., 2018).

## Data availability

The map of the SARS-CoV2 Spike-hACE2 trimer was deposited in the Electron Microscopy Data Bank under EMDB ID 11832 (Spike-ACE2 monomeric complex) and the map of the S1-hACE1 complex under EMDB ID 11833.

## Acknowledgements

We thank Prof. Jason McLellan from University of Texas, Austin for the Plasmids. We thank Lubomir Kovacik and Mohamed Chami for assistance in cryo-electron microscopy. This work was supported by the Swiss National Science Foundation, grant NCCR TransCure. We thank Laurence Durrer and Soraya Quinche from PTPSP, EPFL for work with mammalian cell cultures.

## Author contributions

D.N. did imaging and image processing and data analysis; D.H, K.L. and F.P. produced and purified the protein samples of hACE2 and Spike; F.L. assembled the complex; A.F. performed negative stain TEM. H.S. supervised the project. All the authors edited the manuscript.

## Competing financial interests

The authors declare no competing financial interests.

---

**Figure S1.**
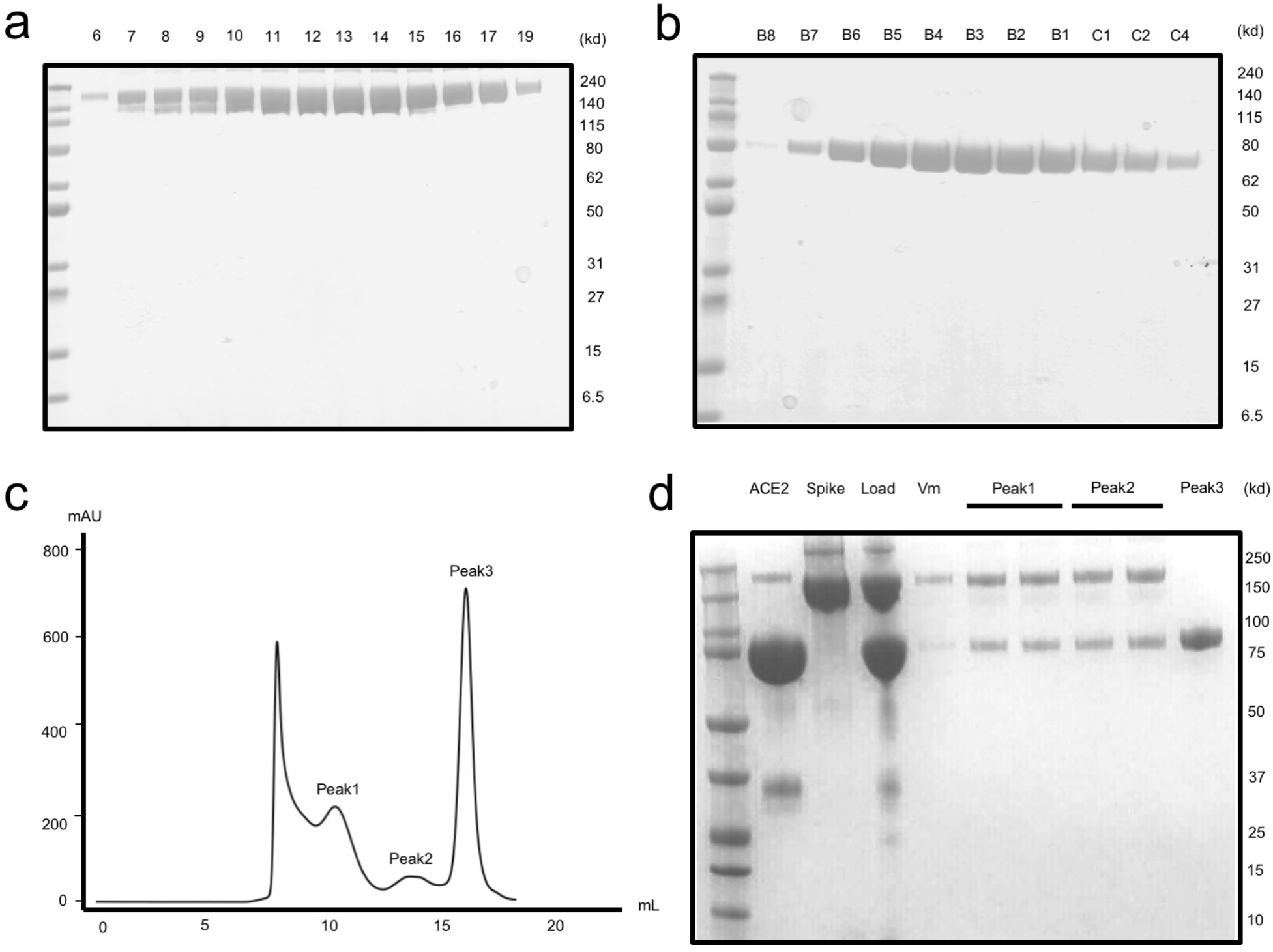
Protein production and purification. **a**. SDS-PAGE of the size exclusion chromatography (SEC) fractions of SARS-CoV-2 Spike (Superose 6 Increase 10/300GL). **b**. SDS-PAGE of the anion exchange chromatography fractions from the purified sample of hACE2 ectodomain (HiTrap Q Sepharose HP). **c**. A representative size exclusion chromatography curve of Spike-hACE2 complex assembly. 280nm UV absorption is shown. **d**. The representative SDS-PAGE of the complex assembly. Lanes hACE2 to Load were after pre-concentration in 100 kDa centrifugal concentrators (Millipore).

**Figure S2.**
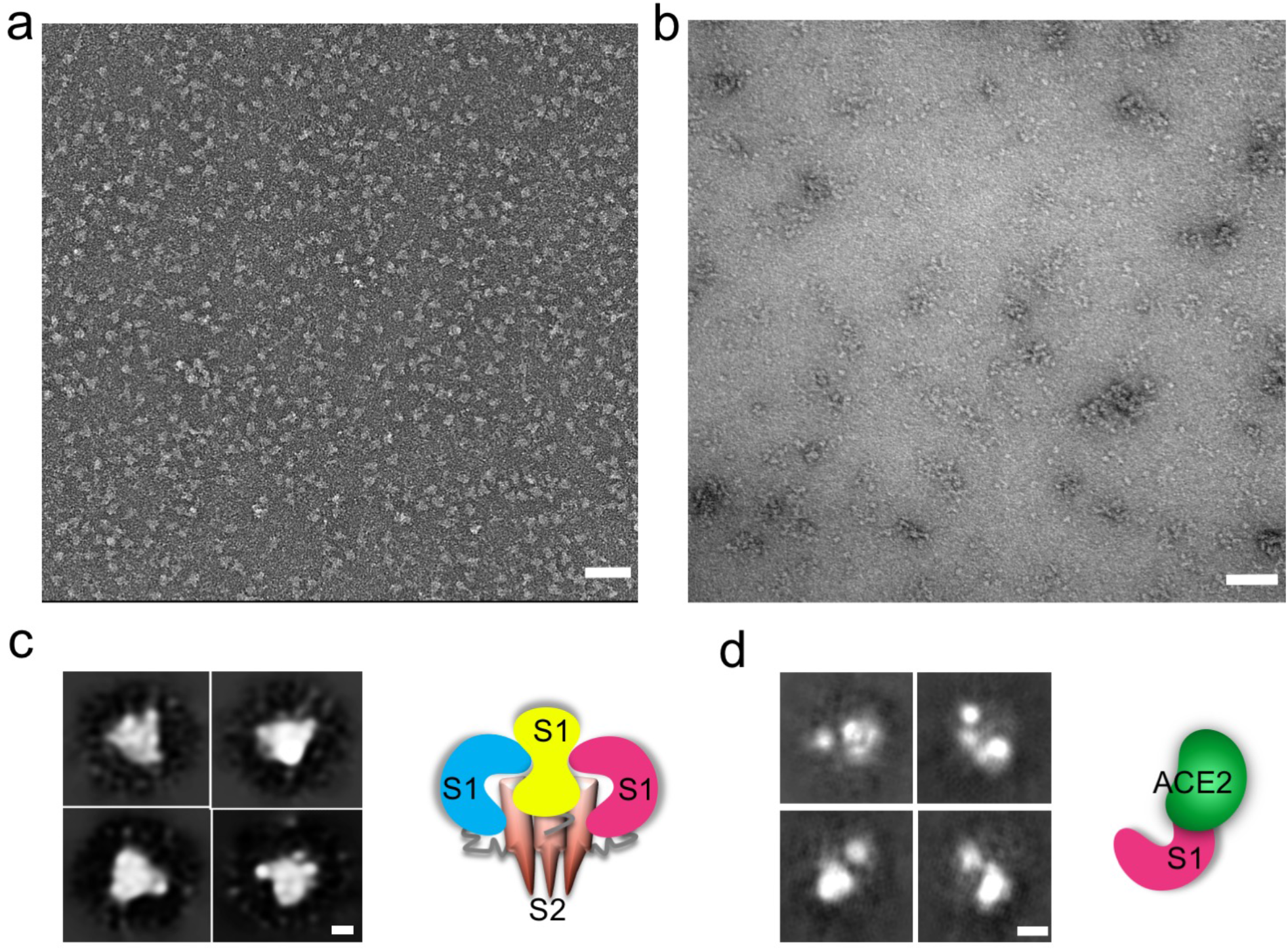
Dissociation of SARS-CoV2 Spike trimer. **a**. Negative stain TEM image of the Spike sample alone (non-incubated sample). **b**. Negative stain TEM image of Spike protein incubated with soluble human ACE2 at a molar ratio of 1:5 for 12 hrs at 4°C, and further purified by SEC. **c**. Left: Representative class averages from particles picked from images as in a. Right: A cartoon interpretation of the structure. **d**. Left: Representative class averages from particles picked from images as in b. Right: A cartoon interpretation of the structure. Scale bars in a and b are 50nm, in c and d are 3 nm.

**Figure S3.**
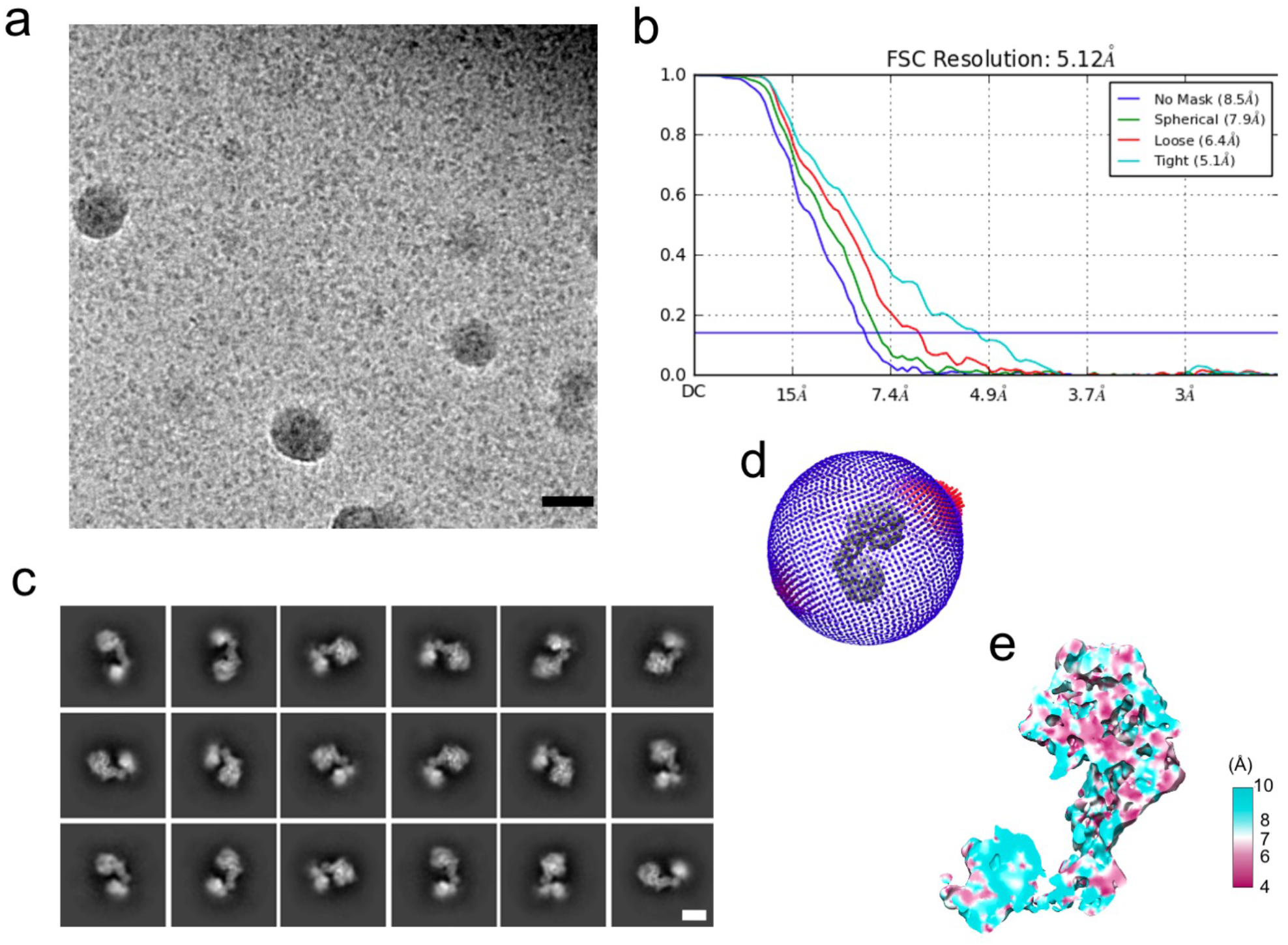
Data quality of Spike-ACE2 monomeric complex (S1-ACE2) sample. **a**. A representative micrograph. **b**. Overall Resolution estimation (FSC, 0.143). **c**. Representative 2D average classes. **d**. Direction distributions. **e**. Local Resolution estimation (MonoRes). Scale bar in a is 50nm and in c is 3nm.

**Figure S4.**
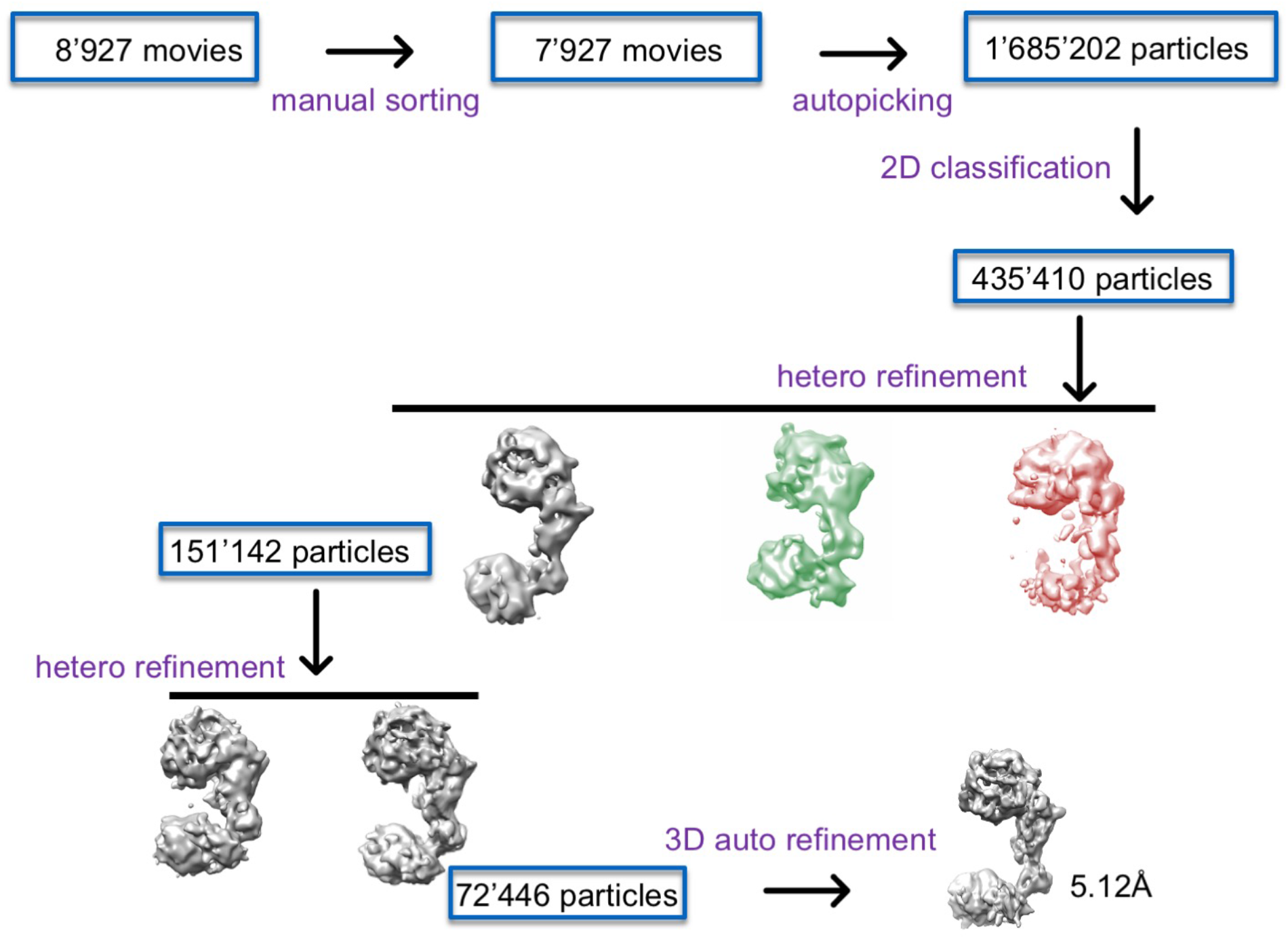
Processing workflow for the Spike-hACE2 monomeric complex (S1-hACE2).

**Figure S5.**
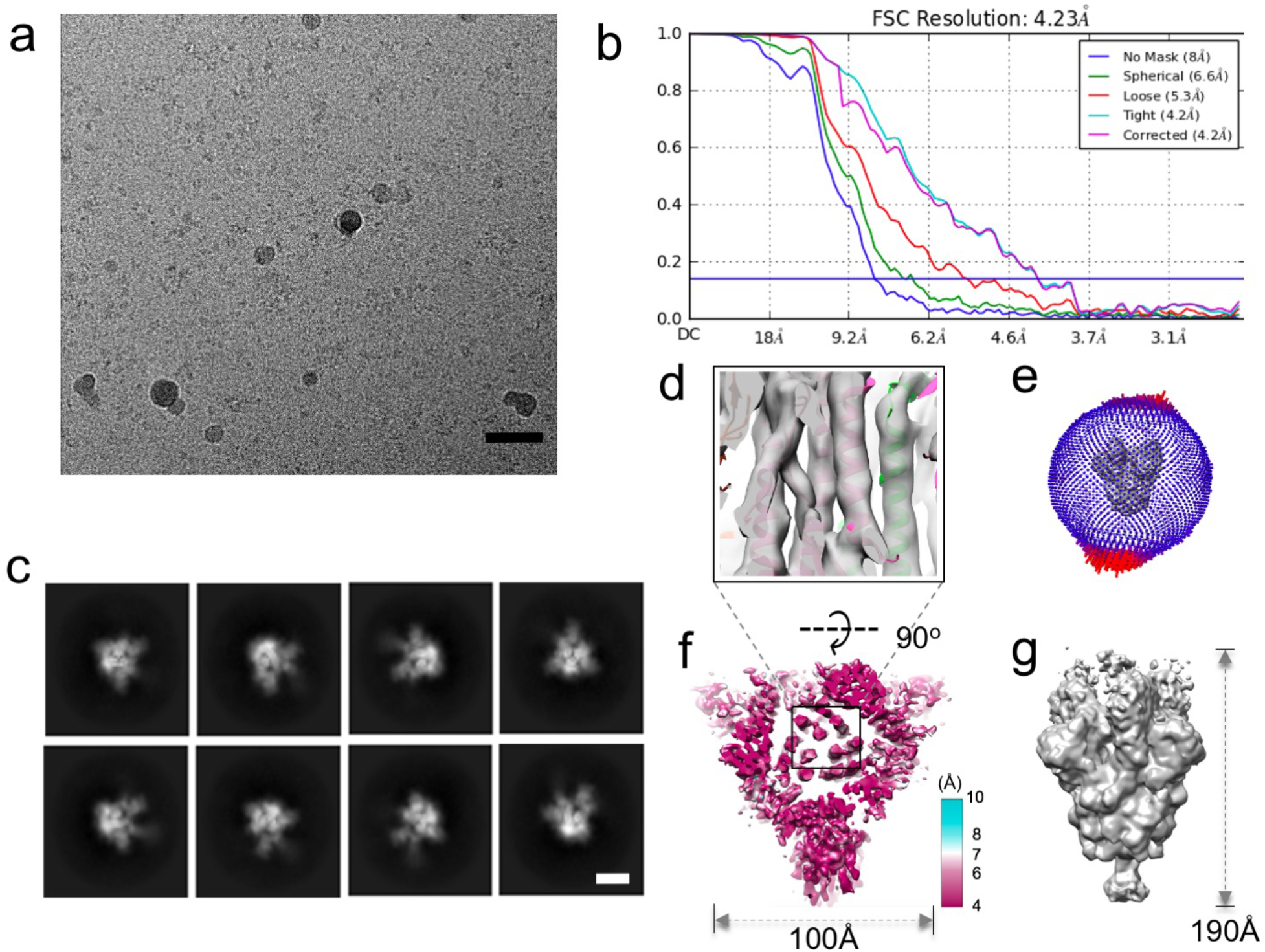
Data quality of the sample of Spike-hACE2 trimeric complex. **a**. A representative micrograph. b. Overall Resolution estimation (FSC, 0.143). c. Representative 2D average classes. d. Model fitted into the S2 trimeric core (The map was low-pass filtered). e. Distribution of particle orientations. f. Local resolution level at the best resolved regions of the trimeric form of Spike-hACE2 complex (bottom view, MonoRes). g. The low-pass filtered EM map at 9Å resolution (For model generation). Scale bar in a is 50nm and in c is 3nm.

**Figure S6.**
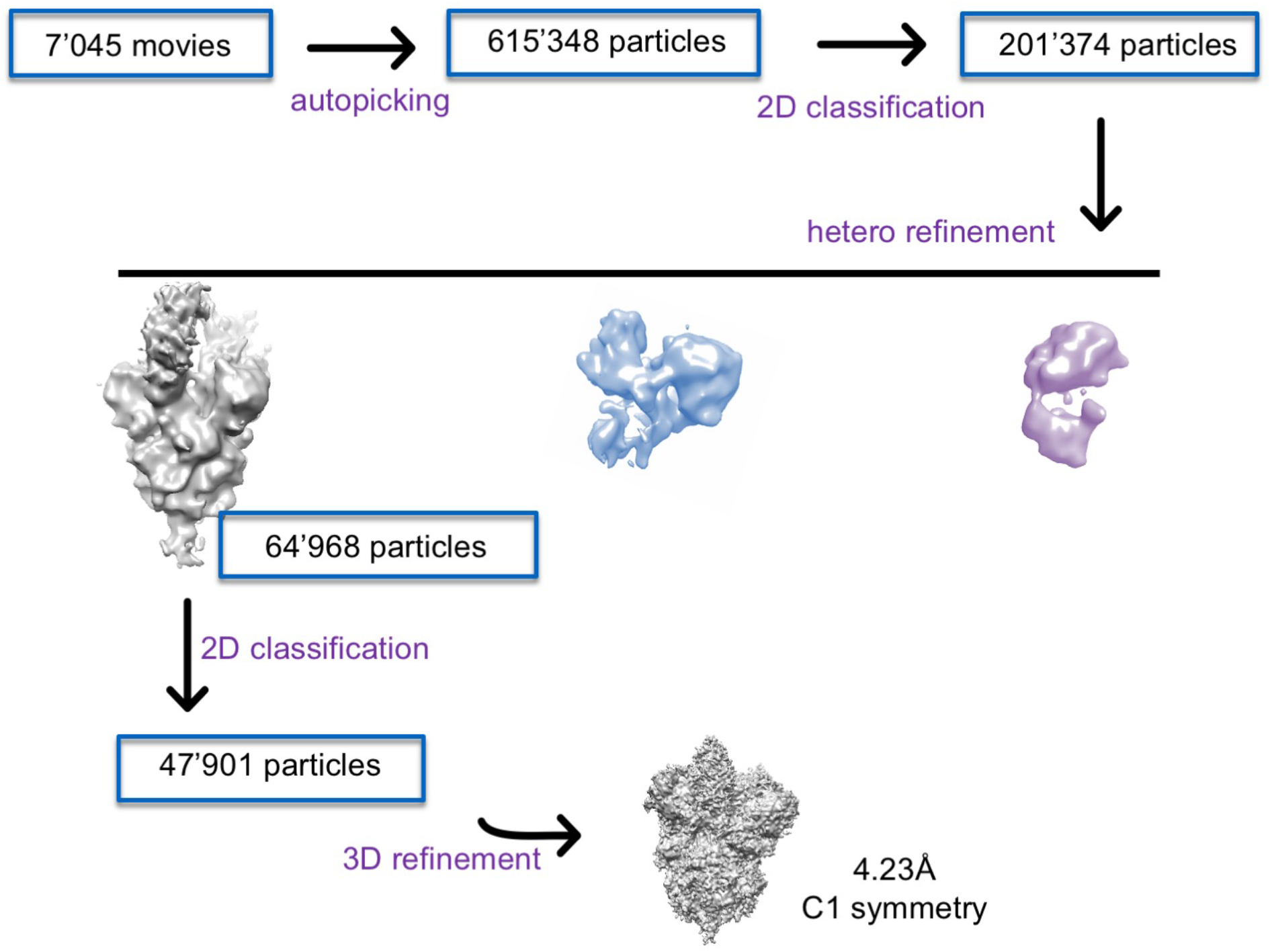
Processing workflows for the trimeric Spike-hACE2 complex.

## Notes

### Competing Interest Statement

The authors have declared no competing interest.

